# On the Digital Psychopharmacology of Valproic Acid in Mice

**DOI:** 10.1101/2020.07.29.220350

**Authors:** John Samuel Bass, Anney Tuo, Linh Ton, Miranda J. Jankovic, Paarth K. Kapadia, Catharina Schirmer, Vaishnav Krishnan

## Abstract

**Objective:** Antiepileptic drugs (AEDs) require daily ingestion for maximal seizure prophylaxis. Adverse psychiatric consequences of AEDs present as: (i) reversible changes in mood, anger, anxiety and/or irritability that often necessitate drug discontinuation, and (ii) autism and/or cognitive/psychomotor developmental delays following fetal exposure. Technical advances in quantifying naturalistic rodent behaviors may provide sensitive preclinical estimates of AED psychiatric tolerability and neuropsychiatric teratogenicity.

**Methods:** Using instrumented home-cage monitoring, we assessed how valproic acid (VPA, dissolved in sweetened drinking water) alters home-cage behavior in adult C57BL/6J mice and in the adult offspring of VPA-exposed breeder pairs. By utilizing a pup open field assay, we also examined how prenatal VPA exposure impacts early spontaneous exploratory behavior.

**Results:** At 500-600mg/kg/d, chronic VPA produced hyperphagia and increased wheel-running without impacting sleep, activity and measures of risk aversion. When applied chronically to breeder pairs of mice, VPA prolonged the latency to viable litters without affecting litter size. Two-week old VPA-exposed pups displayed open field hypoactivity without alterations in thigmotaxis. As adults, prenatal VPA-exposed mice displayed active state fragmentation, hypophagia and increased wheel running, together with subtle alterations in home-cage dyadic behavior.

**Interpretation:** Through automated home-cage assessments of C57BL/6J mice, we capture an ethologically centered psychopharmacological profile of enterally administered VPA that is aligned with human clinical experience. By characterizing the effects of pangestational VPA exposure, we discover novel murine expressions of pervasive neurodevelopment. Incorporating rigorous comprehensive assessments of neuropsychiatric tolerability may inform the design of future AEDs with improved neuropsychiatric safety profiles, both for patients and their offspring.

## Introduction

For the foreseeable future, orally administered antiepileptic drugs (AEDs) ingested daily will remain the first line of defense against seizures in patients with epilepsy. Since AEDs chronically enhance neuronal inhibition and/or diminish excitation without substrate specificity, the high prevalence of anticonvulsant-induced psychiatric and behavioral side effects (PBSEs) should come as no surprise. Such symptoms, which include alterations in mood, anxiety, irritability and anger, substantially contribute to disability in epilepsy^1^, and result in suboptimal dosing, poor compliance and/or discontinuation in up to 25% of patients^2^. Sadly, the highest incidence of PBSEs occur in patients with medically refractory epilepsy and/or pre-existing psychiatric illness^3^, itself associated with seizure intractability^4^. These patients remain at risk for PBSE cross-sensitivity^5^, drastically limiting treatments for a population already vulnerable to suicide^6^ and sudden death^7^. The development of AEDs with improved side effect profiles remains a consensus research benchmark^8^. Nevertheless, preclinical assessments^9^ of AED “tolerability” in rodents remain largely limited to measures of acute motor “toxicity” (e.g., rotarod or open field testing) and/or subjectively scored ordinal scales of well-being (e.g., Irwin screen), which may not capture changes in anxiety, mood, motivation or sociability.

Fortunately, AED-induced PBSEs typically resolve with drug discontinuation^10, 11^. However, in women with epilepsy who are pregnant, AED exposure during fetal neurodevelopment is associated with an increased risk for anatomic and pervasive cognitive/emotional teratogenic effects. Structural malformations (from polydactyly to neural tube defects) may require surgical correction for medical or cosmetic purposes^12^ and may be recognized early (including on prenatal ultrasounds), allowing for early surgical intervention. In contrast, autism, attention deficit and intellectual disability are typically recognized in infancy to early childhood^13, 14^, beyond a theoretical critical period during which medical/behavioral interventions may be preventative. Since clinical trials for new AEDs exclude pregnancy, and since randomized blinded controlled trials of AED safety in pregnancy are unethical, our knowledge of AED teratogenicity has come almost exclusively from pregnancy registries and prospective cohort studies^15, 16^. Broadly, these results confirm that teratogenic side effects are drug-specific and occasionally dose-dependent^17^. Valproic acid (VPA), used as monotherapy or polytherapy is associated with the highest teratogenic risks across both structural and cognitive domains^18–20^. With increased awareness and prescriber education, there have been declines in VPA use during pregnancy and associated declines in malformation rates^21^. However, consensus recommendations regarding the relatively improved neuropsychiatric teratogenic safety profile (*or lack thereof*) for many newer anticonvulsants are conspicuously absent^18, 22^. In support of a direct teratogenic role for valproate, offspring born to pregnant rodent dams (rats or mice) challenged briefly with valproic acid at the approximate time of neural tube closure display behavioral phenotypes deemed “autism-like”, and which are also observed in several genetically informed mouse models of autism^23, 24^. While these studies have unraveled some mechanistically informative insights^25, 26^, two main caveats limit their translational potential. First, anticonvulsant exposure in pregnant women with epilepsy involves dose-adjusted daily intake for the entirety of gestation. Second, abnormalities in “sociability”, repetitive and exploratory behavior are typically deduced from brief “out-of-cage” task-based assays (e.g., three-chamber sociability, open field testing, etc.), which are particularly vulnerable to the confounds of human presence and bias^27–31^.

In this study, we apply instrumented home-cage monitoring^32^ to noninvasively and unobtrusively assess the psychopharmacology of adult and prenatal VPA exposure in C57BL/6J mice. As opposed to VPA injections applied during specific gestational windows^25, 26^ or administered *acutely* to demonstrate *acute* seizure protection^33, 34^, we apply VPA enterally, dissolved in drinking water^35–39^. By combining undisturbed recordings of spontaneous behavior with standardized provocative maneuvers, we illustrate an ethologically sound approach to ascertain drug-induced changes in specific domains of behavior (e.g., feeding, sheltering, etc.) as well as higher-order features of murine behavioral organization^32, 40, 41^.

## Methods

### Mice and Drugs

Experimental protocols were approved by the Baylor College of Medicine Institutional Animal Care and Use Committee (IACUC). All studies were conducted in accordance with United States Public Health Service’s Policy on Humane Care and Use of Laboratory Animals. C57BL/6J mice (#000664, Jackson Laboratories) were bred and weaned at 21 days of age within our vivarium, set to a 12-hour light cycle (lights ON from 0500–1700) under controlled temperature (20–26°C) and humidity (40–70%) settings. Water and chow (PicoLab® Select Rodent 5V5R Standard Diet, with 3.6ppm of folic acid) were provided *ad libitum*. Cages were furnished with cellulose bedding (Biofresh). For VPA dosing, we differentiate between *intended* doses and *actual doses* (determined by bottle and body weight measurements). In pilot experiments with 6-10 week-old C57BL/6J mice, actual doses of ~600mg/kg/d were obtained by dissolving 2.6mg/ml of sodium valproate (Sigma Aldrich) in drinking water (sweetened with 0.8% sucrose). Pentylenetetrazole (PTZ) was dissolved in sterile-filtered normal saline and injected intraperitoneally (30mg/kg, 5ml/kg). To model fetal exposure, breeder pairs were assembled and received VPA or control solutions that were changed weekly and replaced with standard drinking water at parturition, determined by daily external cage checks. At the end of experiments, mice were euthanized by CO_2_ inhalation.

### Home-cage monitoring and video-tracking

Remote behavior monitoring was conducted as previously described^32^ within a satellite facility termed the BMU (behavior monitoring unit) replicating vivarium lighting, light cycle, humidity and temperature settings. Mice were housed within one of sixteen Phenotyper (Noldus Information Technology) home-cages (30×30×47cm) with clear plastic walls, one or two water sources fitted with lickometers (detecting capacitance changes), a feeding meter (measuring beam breaks) and a detachable running wheel (utilizing a wheel-attached magnet and a wall-attached magnet sensor). Bedding and chow were identical to those provided in vivarium cages. A sound machine (‘Lectrofan, Adaptive Sound Technologies) played continuous white noise. An aerial video-feed was established using a ceiling-mounted infrared camera together with an array of ceiling mounted infrared lamps. An infrared-lucent shelter (10×10×5-6cm) was deployed for all non-dyad studies. The “shelter zone” was defined on a distance-calibrated arena for each cage. Recordings were initiated at a specific clock time or started manually (Fig. 2K, 4B, 5B). Live center-point tracking was conducted via Ethovision XT 14 (Noldus) using dynamic subtraction at a sample rate of 15/s. Licking, feeding and wheel-running data were integrated through a hardware control module. For light spot testing^32, 42^ (always performed between 1900-2000), a ceiling-mounted white LED produced a gradient of light, measuring ~670 lux at the left upper quadrant and ~27 lux at the right lower quadrant of the cage. A 60s long 2300Hz “beep” was then administered at 2100. To avoid phenotypic drifts secondary to the experience of long-term voluntary exercise^43^, running wheels were affixed and subsequently detached within 20-23h. Convulsions were detected by integrating video and real-time changes in mobility, as described previously^32^. Dyads (in Fig. 5) were assembled as sex- and condition-matched pairs of singly-housed mice (depicted in Fig. 4). Since subjects were unmarked, tracking results are presented as “sum distances” [of both mice] and distance between subjects (DBS). Satellite access was restricted to CS, JSB and VK who wore a gown, cap, face mask and gloves at all times, and entered at least once daily to visually inspect food and water sources, assess general mouse wellbeing and/or administer injections. Satellite veterinary inspections were performed every other week (~1100–1200).

### Pup Open Field

Pup video recordings were conducted within a square open field made of Lightaling building bricks (Amazon) with a side 14.5cm and 7.7cm high walls. This brick frame was positioned over an absorbent bench pad warmed over a circulating water blanket and replaced after every set of mice studied. A KF-TM2515T tripod (K&F Concept) combined with a Kimire Digital Camcorder (1080P, 24 megapixels) was used to obtain 15-minute long aerial open field recordings. Ethovision XT was configured to identify time in “center” (a concentric square of 7.25cm side) and entries into each of four “corners” (3.6 x 3.6cm). Serial measurements of open field behavior (without drug exposure, Fig. 3D) were performed on a large cohort of C57BL/6J pups, for which mice were subjected to no more than three evaluations separated by at least 3-4 days. In contrast, for Fig. 3E, all pups were studied *serially* at P5, P10 and P15 (post-natal day 15).

### Data analysis

Graphs were plotted and analyzed with Prism GraphPad 8 and Microsoft Excel 2016 (Fig. S2 C, D). Time budgets (e.g., Fig. 1C) were calculated by tallying total *durations* of sheltering, feeding and drinking. “Sleep” epochs were computed as at least 40s-long contiguous time periods devoid of “movement” (defined as sample velocity ≥ 1.2cm/s), an approach that has been previously validated electroencephalographically^44^ and pharmacologically^32^. Active states were defined as epochs lasting at least 1 minute long during which mice consistently traversed at least 5cm/min (see Fig. S2 for illustration). For dyads, social bouts were defined as at least 3s-long epochs during which DBS consistently measured ≤ 4cm. Unpaired two-tailed Student’s *t* tests were used to compare two group means. Repeated measures analysis of variance (RMANOVA) was conducted by fitting a mixed effects model, with main effects and interactions listed in Table S1. Two-tailed Chi squared analyses were applied to compare categorical variables across two groups.

**FIGURE 1:**
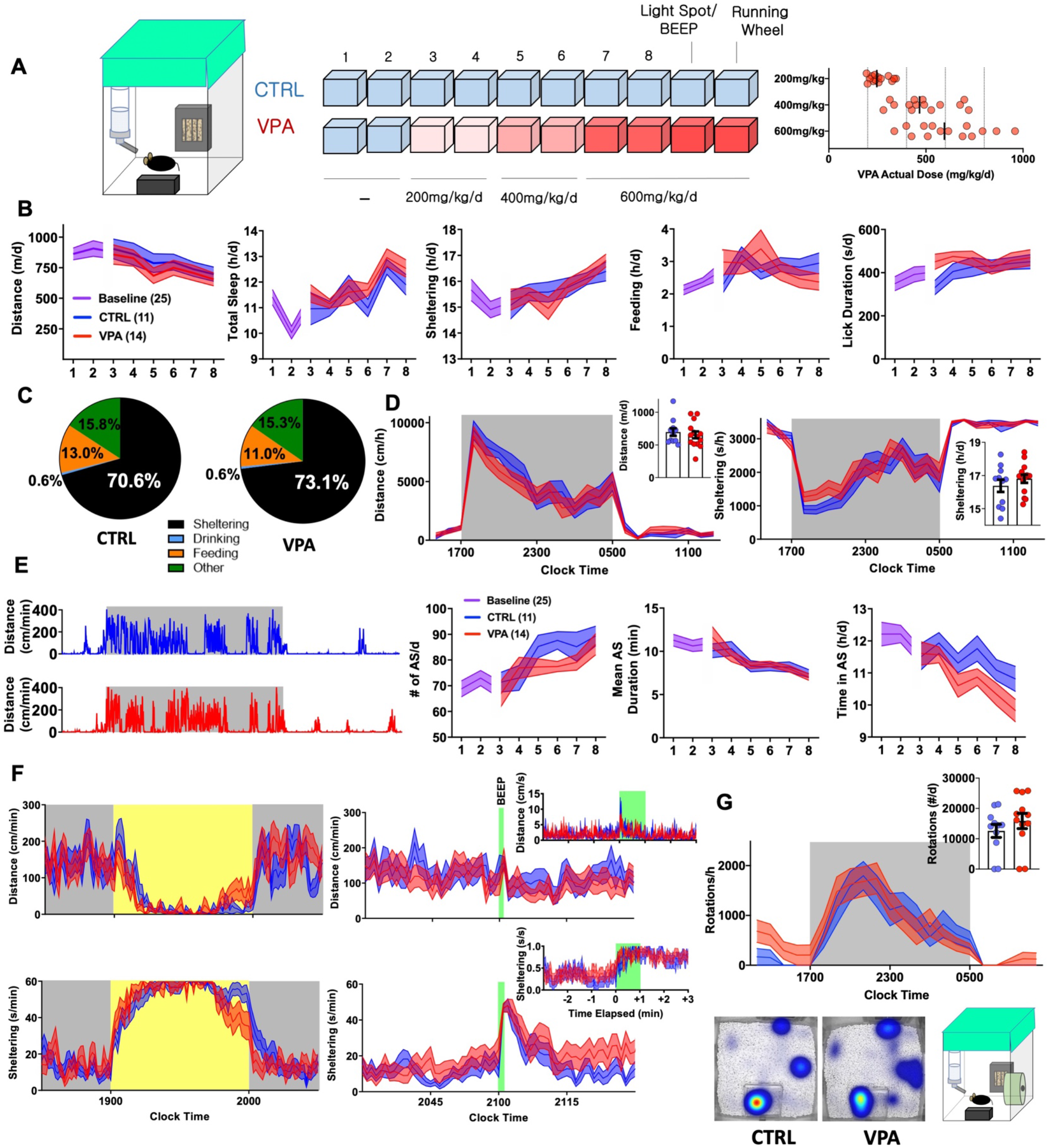
Early Behavioral Consequences of VPA Exposure (CTRL n=11 [5 female] VPA n=14 [7 female]). (A) Schematic of 10-day home-cage protocol, during which VPA is increased to an intended daily dose of 600mg/kg/d. RIGHT: Actual consumed daily doses (derived from bottle and body weights). (B) Home-cage behavioral parameters represented as daily totals (see table S1 for repeated measures analyses). (C) Time budgets, and (D) hourly/total distances and sheltering parameters measured on day 8. (E) Representative actograms (day 8) and quantification of active state parameters. (F) Effects of light spot (left) and “beep” testing (right) on measures of distance and sheltering (day 9). (G) Wheel-running (day 10) with representative heat maps of position probability. Mean ± S.E.M shown for all. CTRL: control, VPA: valproic acid, d:day.

## Results

### On the Initiation of Valproic Acid Therapy

VPA is pharmacodynamically complex (inhibiting various ion channels, GABA degradation enzymes and histone deacetylases) and displays complex pharmacokinetics, with significant protein binding and biologically active metabolites with varied brain clearance rates^33, 45^. As an anticonvulsant, VPA dose requirements vary widely, resulting in poor correlations between plasma concentrations and anticonvulsant efficacy^33^. Thus, rather than aim for a specific plasma or brain concentration, we began by exploring the effects of VPA dissolved in drinking water (*o.s.*) at a dose range of ~500-600mg/kg/d, previously demonstrated to provide some seizure protection in rodent models^35–38^. 8-9week old C57BL/6J mice were admitted to BMU home-cage chambers fitted with a single lickometered water source (0.8% sucrose drinking water), an infrared-lucent shelter and food hopper (fitted to sense beam breaks). Following two 23h-long baseline recordings, mice were randomized to receive treatment with either control or VPA solutions at intended doses of 200 (days 3,4), 400 (days 5,6) and 600mg/kg/d (days 7, 8, Fig. 1A). We hypothesized that behavioral parameters would remain relatively stable in controls, while VPA-treated mice would display dose-dependent changes in activity and/or neurovegetative function. Surprisingly, *both* groups displayed downward drifts in daily total distances accompanied by gradual increases in sheltering and “sleep” (Fig. 1B, computed noninvasively^32^, see Methods). Stepwise increases in VPA dose were not associated with evidence of acute taste aversion, as assessed by lickometry (Fig. S1). On day 8, VPA-exposed mice consumed an average actual dose of 616mg/kg/d (range: 337-959), which *positively* correlated with licks/day (r=+0.59, p<0.05) *and* total daily distances (r=+0.46, p=0.09, n=14), the latter hinting towards dose-dependent *activation* rather than sedation. Compared with controls, VPA-treated mice displayed a modified average time budget^32^, accumulating greater total shelter time at the cost of feeding. Overall, hourly measures of total distance and sheltering were largely similar between groups (Fig. 1C, D).

To resolve finer changes in activity patterns, we visualized daily actograms resolved to cm/min (Fig. 1E). In natural environments, animals oscillate between *active* (patrolling, foraging, grooming) and *inactive* states (rest, sleep or quiet wakefulness). In mice, active state structure varies exquisitely by background strain^40^ and is modulated by environmental and genetic factors, including alterations in energy balance^41^. Unlike states of consciousness (e.g., wakefulness or sleep) that are formally defined electroencephalographically, active states can be discerned through quantitative and continuous measures of behavioral output. We defined active states as at least 1-minute long epochs displaying a sustained mean velocity of 5cm/min (Fig. S2). While active states in both groups evolved to become more frequent and shorter in duration, higher doses of VPA produced a deficit in total active state time (Fig. 1E). Over the next two days at the same dose, we applied a series of provocative maneuvers. During a light spot test^32, 42^, which imposes a conflict between nocturnal foraging behavior and light avoidance, VPA-treated mice displayed similar patterns of hypoactivity and shelter entry, but prematurely emerged from their shelters towards the stimulus end. An hour later, a 60s long “beep” produced a transient activity spike and shelter entry that was similar between control and VPA-treated mice (Fig. 1F). To obtain a final measure of wellbeing, we affixed running wheels within their home-cages. Compared with controls, VPA-treated mice displayed increased early interest in wheel-running and similarly robust rates of nocturnal running (Fig. 1G). Together, these results depict the early behavioral response to *o.s.* VPA therapy, characterized by slight activity reductions without evidence of neophobia, altered sensory processing or diminished exercise motivation.

### On Ethograms of Chronic VPA Exposure

To profile the tolerability of chronic *o.s.* VPA, group-housed 5-6-week old C57BL/6J mice were exposed to either VPA or control solution for four weeks within our vivarium (Fig. 2A), during which VPA-treated mice consumed approximately 500-700mg/kg/d without altering weight gain (Fig. 2B, C). Mice were then transferred to BMU home-cages fitted with *two* lickometered water sources (water vs 0.8% sucrose), and VPA exposure was continued (for VPA-treated mice only) in *both* bottles. On day 2, we observed no changes in total daily distances or “sleep”, but VPA-treated mice displayed significantly greater feeding behavior and trended to display lower sucrose preference (p=0.07, Fig. 2D-F). Chronic VPA did not impart changes in active state frequency or mean duration, and both groups displayed similar responses to light spot and “beep” provocations (Fig 2G-I). While lowered sucrose preference may suggest anhedonia, VPA-treated mice displayed a marked increase in wheel-running and as a group, displayed significantly fewer non-runners^30^ (χ^2^ = 9.62, p<0.01). Finally, all mice received a daytime (~1200) intraperitoneal injection of pentylenetetrazole (PTZ)^33, 34, 46^. In identical home-cages, we have previously shown that subconvulsant dose PTZ injections in C57BL/6J mice acutely produce immobility together with deficits in sheltering. Repeated PTZ injections (kindling) improve latencies to shelter re-entry, albeit at the expense of increased convulsion likelihood^32^. In this light, VPA ingestion produced a behavioral profile resembling *early* PTZ injections, with fewer tonic-clonic convulsions (χ^2^ = 2.39, p=0.1) but more profoundly impaired sheltering (Fig. 2K). Thus, at doses that modify PTZ responses, chronic *o.s.* VPA produces hyperphagia and increased voluntary wheel-running without significantly altering sensorium, sleep or active state structure.

**FIGURE 2:**
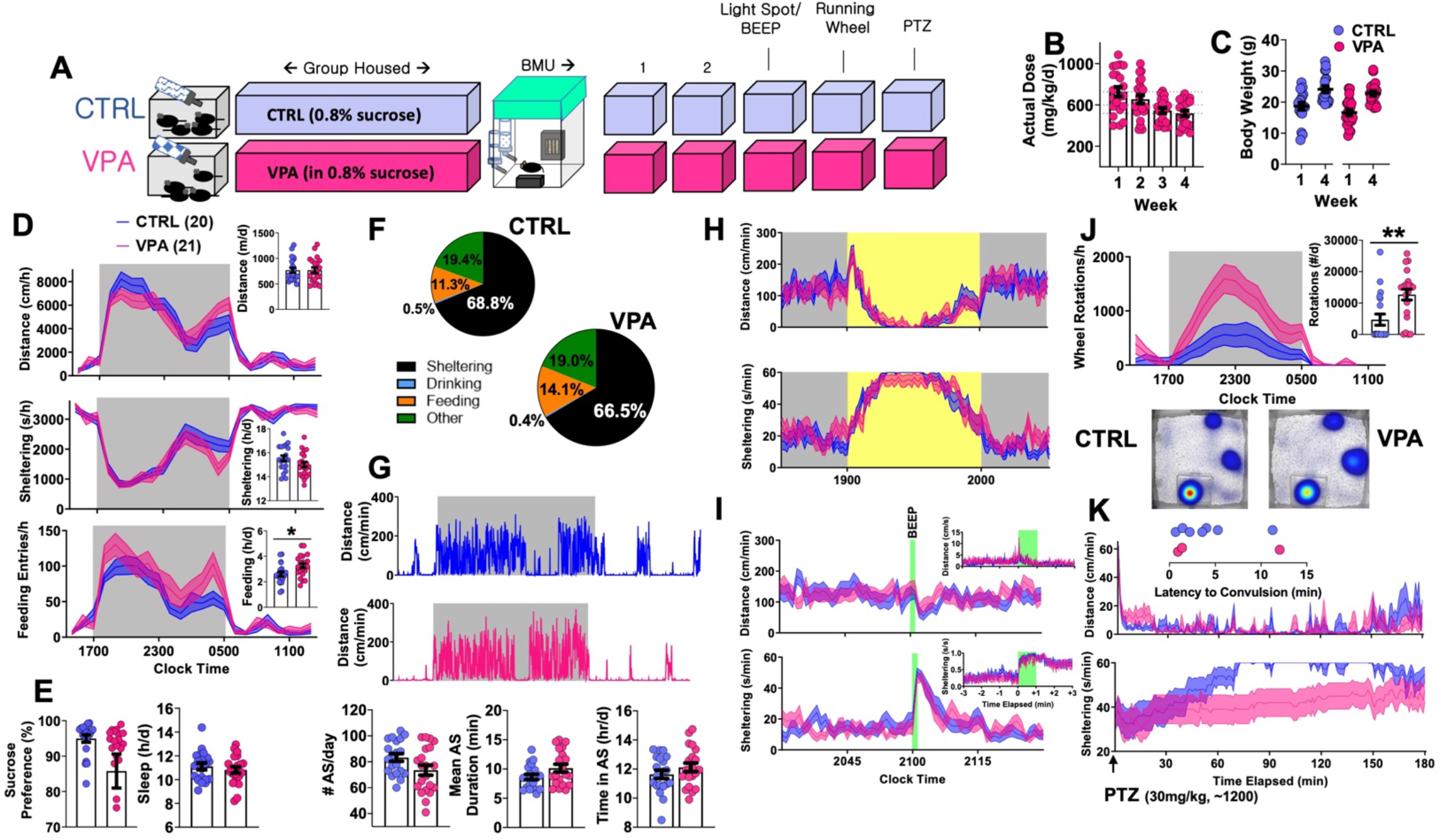
Behavioral Consequences of Chronic VPA Exposure (CTRL n =20 [11 female], VPA n=21 [11 female]. (A) Schematic of protocol, with four weeks of VPA exposure at an intended daily dose of 600mg/kg/d, followed by five days of BMU recordings. (B) Estimates of actual consumed daily doses. (C) Weight change over four-week treatment period. (D) On day 2, total and hourly measures of distance, sheltering and feeding, presented as both feeding duration and feeding entries. (E) Sucrose preference, behaviorally defined “sleep” and (F) time budgets all obtained from day 2. (G) Representative actograms and quantification of active state parameters (day 2). (H) Effects of light spot and (I) “beep” testing on measures of distance and sheltering. (J) Wheel-running behavior with representative heat maps of position probability. (K) Changes in distance and sheltering following a single 30mg/kg intraperitoneal PTZ injection, with inset depicting latency to convulsive seizures. Mean ± S.E.M shown for all. CTRL: control, VPA: valproic acid.

### On the Early Consequences of Pangestational VPA Exposure

To measure neuropsychiatric teratogenicity, 8-9-week old sexually naïve C57BL/6J breeder pairs were assembled and randomized to receive either control solution or VPA. At parturition, bottles were replaced with sucrose-free drinking water (Fig. 3A). VPA exposure in this manner prolonged the latency to a viable litter by ~2 weeks, without impacting litter size (Fig. 3B) or sex ratios (control: 57% female, VPA: 48% female, p>0.1). Externally obvious sequelae of neural tube maldevelopment or other structural anomalies were not observed in either group. We then sought to examine how pangestational VPA exposure impacted early neuropsychiatric development without imposing prolonged maternal separation. Most available tests to assess early murine neurological milestones emphasize sensorimotor reflexes (e.g., cliff avoidance, negative geotaxis) and are subjectively scored on ordinal scales^47, 48^. To achieve unbiased quantitative measurements of pup behavior suitable for automated analysis, we devised a “pup open field” and conducted 15-minute long aerial video recordings that were subsequently video-tracked. We first examined a large cohort of unmanipulated C57BL/6J pups assessed between P5 (postnatal day 5) and P15. Total open field distances remained stably low until ~P12, corresponding to the timing of eye opening^48^. In contrast, open field center times displayed two stepwise decrements at P9 and P12, illustrating how thigmotaxis emerges prior to eye opening (Fig. 3C, D). Control and VPA-exposed pups behaved similarly at P5 and P10. At P15, VPA-exposed pups displayed significantly lower total open field distances without changes in center time or maximum velocity (Fig. 3E, F). Heat maps of position probability identified large individual variations in field exploration across both groups (Fig. 3G). VPA-exposed pups were less likely to explore all four corners of the open field (χ^2^ = 9.62, p<0.05), collectively illustrating a deficit in early measures of spontaneous exploratory behavior.

**FIGURE 3:**
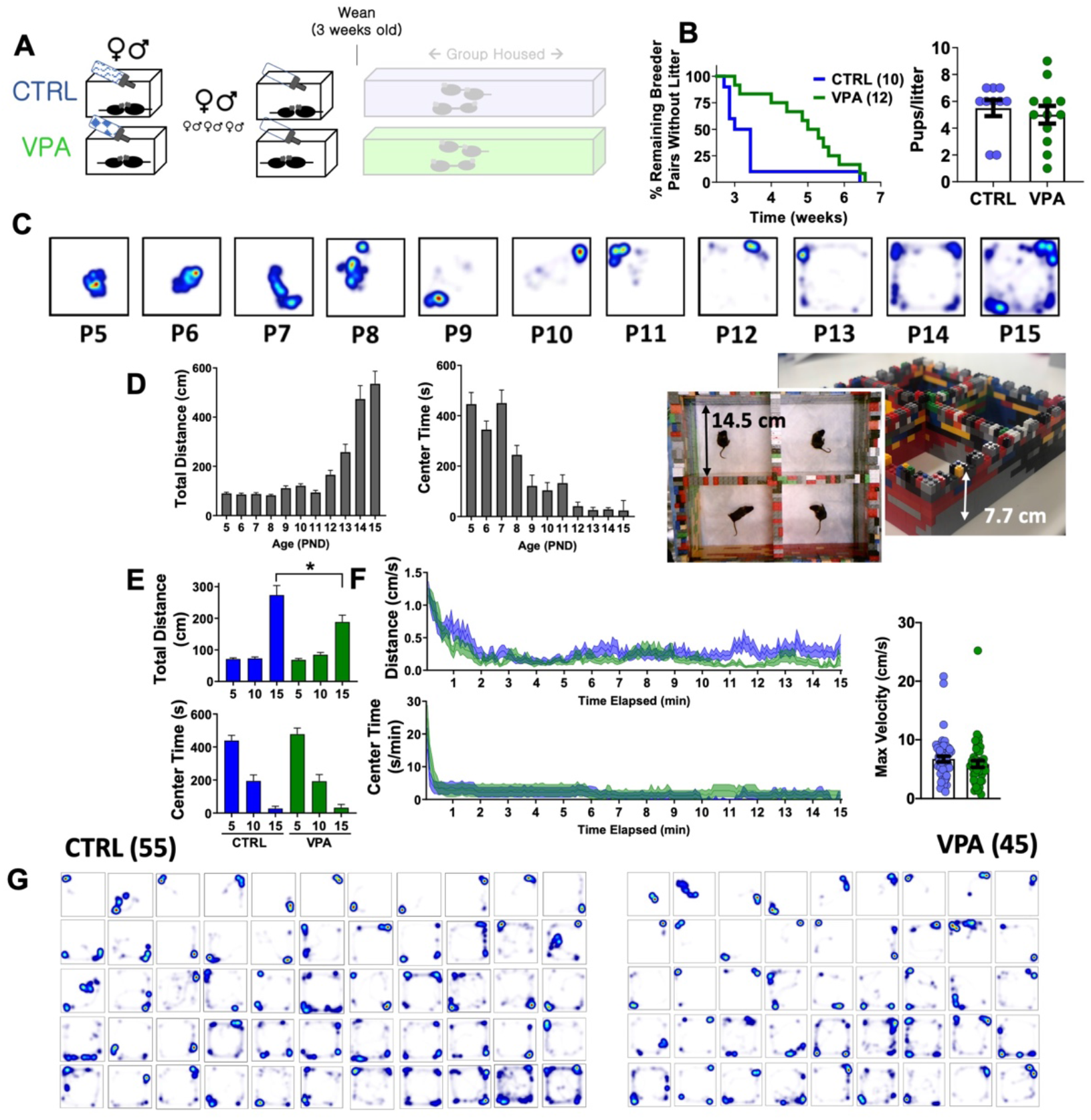
Pup open field behavior following prenatal VPA exposure. (A) Breeder pairs were exposed to VPA or control solutions until parturition. (B) VPA-exposed breeder pairs took longer to produce a viable litter (p<0.05, Mantel-Cox log-rank test), without a change in litter size. (C) Representative heat maps of position probability from a separate cohort of C57BL/6J mice (n=27—45/group), illustrating progressive changes in open field exploration. (D) Distances and center times for C57BL/6J mice. (E) Total distances and center times across CTRL and VPA cohorts, and (F) time-dependent changes in behavior at P15. (G) Heat maps of position probability for all pups studied at P15 (CTRL n = 55 [32 female], VPA n=45 [21 female]). Mean ± S.E.M shown for all. CTRL: control, VPA: valproic acid. P15 = post-natal day 15.

### On the Effects of Pangestational VPA Exposure in Adulthood

When first introduced to BMU home-cages at ~8-9weeks of age, prenatal-VPA exposed adult mice displayed a similar pattern of locomotor habituation with slightly greater total sheltering times (p=0.1, Fig. 4B). On their second baseline day, VPA-exposed mice displayed similar rates of overall movement, sleep and sucrose preference. Daily time budgets revealed a relative increase in sheltering behavior at the cost of feeding and “other” activities (time spent *not* feeding, drinking or sheltering, Fig. 4C, D). VPA-exposed mice displayed a significantly greater number of daily active states (~81 vs 71/day in controls) that were of significantly shorter duration (~10.7 mins vs 9 mins), and a reduced total active state time (Fig. 4E). Light spot and “beep” tests revealed no significant group differences (Fig. 4F). When presented with a running wheel, VPA-exposed mice displayed significantly greater average rates of nocturnal wheel running (Fig. 4G).

**FIGURE 4:**
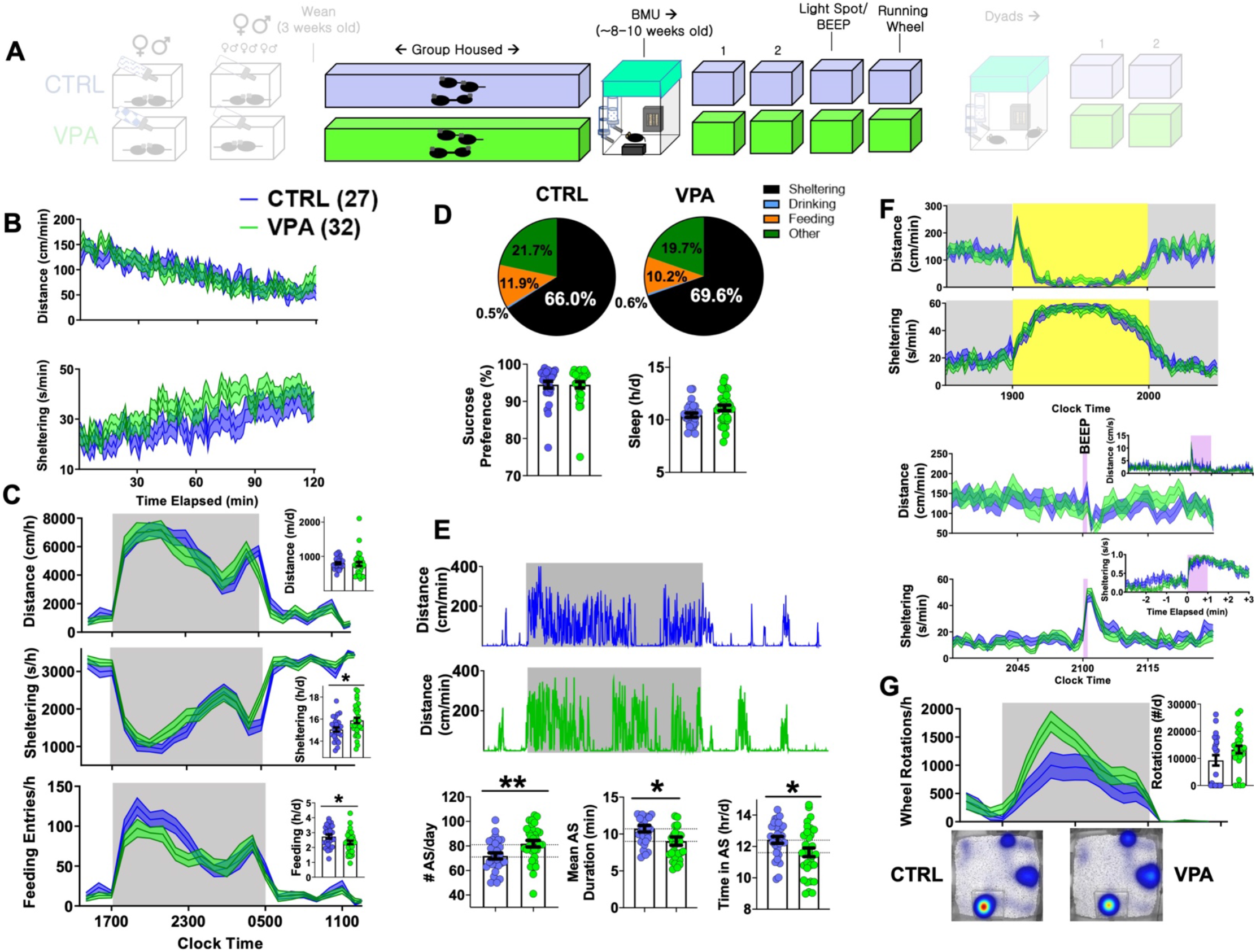
Adult home-cage behavioral profile following prenatal VPA exposure. CTRL n=27 [14 female], VPA n=32 [14 female]. (A) After weaning, mice were group housed until 8-9 weeks of age. (B) Dynamic changes in locomotor activity and sheltering behavior in the first two hours of BMU recording. (C) Hourly/total measures of distance, sheltering and feeding (day 2), and (D) corresponding time budgets, sucrose preference and measures of “sleep”. (E) Representative actograms and quantification of active state parameters. (F) Effects of light spot and “beep” testing on measures of distance and sheltering. (G) Wheel-running behavior with representative heat maps of position probability. Mean ± S.E.M shown for all. CTRL: control, VPA: valproic acid.

Autistic endophenotypes of the human fetal VPA syndrome^13–15, 22^ have been ascertained in rodent prenatal VPA protocols using tasks of sociability, social novelty and/or ultrasonic vocalizations^23, 25^. To examine how prenatal VPA exposure altered the dynamics of social behavior within a now familiar home-cage setting, we next paired sex- and drug exposure-matched mice into “dyads” (CTRL-CTRL vs VPA-VPA, Fig. 5A). Rather than manually score specific social behaviors like sniffing or following^24^, we simultaneously tracked both mice to capture kinematic measures and a continuous readout of inter-mouse proximity, termed DBS (distance between subjects). In the first two hours, mean DBS levels reduced from ~15cm to ~10cm in both groups, paralleling a locomotor habituation response (Fig. 5B). We then closely examined the morphology of individual social bouts, which we defined as epochs lasting ≥ 3s where DBS was consistently ≤ 4cm. Prenatal VPA exposure did not impact the frequency or the mean duration of social bouts (Fig. 5B). Over the subsequent two days, circadian alterations in mean proximity generally mirrored locomotor fluctuations^49^. Prenatal VPA-exposed dyads displayed a delayed dark-light transition response^50^, but were nevertheless noted to huddle together during the day time (Fig. 5C). Social bouts were generally brief during the dark phase, but during a light spot challenge, VPA-exposed mice displayed significantly longer social bout durations (Fig. 5D). “Beep” testing did not elicit differences in dyadic responses (Fig. 5E). Finally, to interrogate seizure threshold, we applied subconvulsant dose PTZ injections to all mice. While convulsions occurred at a low rate in both groups (χ^2^ = 0.75, p>0.3), continuous objectively derived home-cage measures were distinct: VPA-exposed mice displayed a blunted motor recovery and remained more distant from their counterparts (Fig. 5F), suggesting enhanced seizure severity^51, 52^.

**FIGURE 5:**
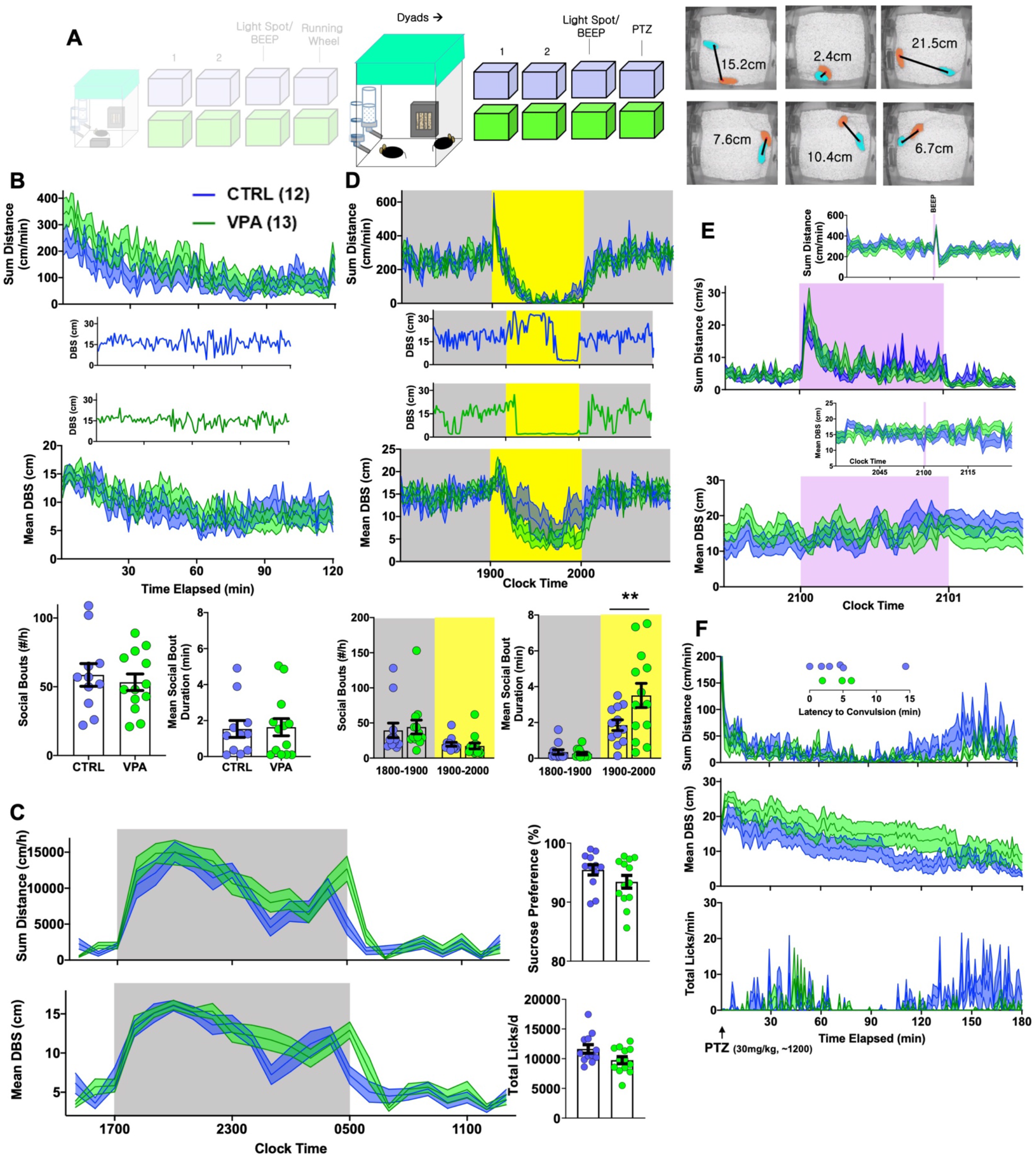
Effects of prenatal VPA exposure on dyadic behavior. (A) Sex- and group-matched dyads were paired within BMU home cages. Videotracking was used to calculate proximity (distance between subjects/DBS). (B) Continuous measures of total distance and DBS during the initial two hours of this encounter, with representative examples of DBS. BOTTOM: Total social bouts and mean social bout durations. (C) Hourly measures of total distance and mean DBS, with average sucrose preference and total licks (day 2). (D) Light spot testing revealed greater social proximity in prenatal VPA-exposed dyads, and results of representative dyads are shown. (E) “Beep” test and (F) dynamic changes in distance, DBS and licking behavior following a 30mg/kg PTZ injection applied to both members of the dyad, with inset depicting latency to convulsive seizures. Mean ± S.E.M shown for all. CTRL: control, VPA: valproic acid.

## Discussion

In the 1960s, after the serendipitous discovery of VPA’s anticonvulsant actions (against PTZ in rodents), the earliest human trials of VPA were conducted in psychiatric asylums, often enriched with epileptic patients^53^. Valproate displayed both anticonvulsant *and* beneficial psychotropic properties, with improvements noted in depression, “viscosity” and occasionally mild euphoria^53, 54^. Following its release in the US in 1978^55^, VPA experienced enormous popularity both as a broad-spectrum anticonvulsant^56^ and mood stabilizer^53^, and our knowledge of VPA’s systemic adverse effects expanded, including pancreatic and hepatic toxicity, platelet dysfunction, weight gain, sexual dysfunction^57^ and tremor^58^. A thorough understanding of the structural and cognitive consequences of fetal VPA exposure did not emerge until the 1990s-2000s, a rather extended and potentially preventable delay that contributed to significant adverse developmental outcomes^59^. Rodent studies on VPA’s psychopharmacological effects lagged well behind human clinical experience, with demonstrations of anxiolytic-like effects (1985)^60^, antidepressant-like effects (1988)^61^, and evidence of “autism-like” behavior in rodents prenatally exposed to VPA on embryonic day 12.5 (2005)^62^.

Today’s newest generation of anticonvulsants are intelligently designed through targeted drug discovery programs that combine the latest technologies in high-throughput screening and medicinal chemistry with an array of established anticonvulsant assays^9, 63, 64^. Still, evaluations of tolerability crudely focus on motor *toxicity*, which itself may poorly correlate with complaints of somnolence or fatigue^65^. Similarly, animal studies on teratogenicity focus primarily on structural malformations, and standardized assessments of neurodevelopmental behavioral toxicity have been consistently deferred and/or ignored^66^. In this study, we asked whether detailed automated evaluations of home-cage behavior would be sensitive enough to detect anticonvulsant psychotropic effects, and further, reveal features of pervasive neurodevelopment (if any) in mice exposed to that anticonvulsant prenatally. We chose VPA as a prototypical AED known to display both such features, and employed C57BL/6J mice as subjects, given their widely reproducible home-cage ethograms^32, 50, 67^ and low rates of within-strain variability^68^. Home-cage monitoring is grounded on the assumption that *if* a rodent were to experience clinically meaningful symptoms of autism, developmental delay, anxiety or depression, such a syndrome would exert measurable effects on activities of daily living as measured in *their* home-cages. Extended recording durations highlight phenotypes which may be only evident during a specific circadian phase. This is commonly observed in actigraphic evaluations of patients with psychiatric disorders, where psychomotor changes are largely limited to the patient’s biological day^69, 70^. In view of this point, home-cage monitoring is unparalleled in its ability to characterize the rich dynamic behavioral features of the murine “night”.

We devised two paradigms to assess the tolerability of VPA ingestion. First, we examined how mice responded to the initial introduction of VPA during a gradual dose uptitration schedule, akin to a within-subject “placebo-controlled” dose-response study. During daily recordings conducted over 8 days, control mice displayed downward drifts in overall cage exploration, with increases in sheltering, feeding and “sleep”. Active states also evolved to become more frequent and shorter in duration in control mice. These findings may relate to infradian rhythms^71^ and/or represent an *observer* effect: by *individually* housing our mice (albeit within an enriched home-cage) to *individually* assess changes in behavior, we observed phenotypic drifts that may be secondary to the cumulative effects of social isolation. Among various trajectories measured, VPA at doses of 400-600mg/kg/d diminished total active state time without significantly altering the mean duration of active states. We infer that VPA, gradually introduced in this manner, made mice somewhat “less active”. Nevertheless, VPA consumption did not alter voluntary wheel-running behavior, a task that is broadly sensitive to deficits in motivation, neuromuscular function and/or neophobia^30^.

In the second approach, we applied similar doses of VPA chronically to group-housed mice. While we did not observe changes in weight gain over the treatment period, VPA-treated mice accumulated significantly greater feeding durations and entries without clear evidence of locomotor suppression, active state dysregulation or increased “sleep”. Weight gain is a well-known side effect of VPA and may be explained by increased appetite, although appetite, satiety or hunger are challenging to objectively measure in humans^72^. VPA has been linked to hyperinsulinemia (and insulin resistance), and hyperleptinemia (with leptin resistance), both of which would be expected to increase appetite^58, 73^. A mild reduction in sucrose preference was observed, consistent with a state of lower natural reward sensitivity (*anhedonia*). Nevertheless, VPA-treated mice displayed a robust increase in overall wheel-running, which may be interpreted as adaptive and beneficial (antidepressant-like, pro-resilient) or maladaptive and pathological (pro-manic, “compulsive/addiction-prone”)^30^. Ultimately, these data emblemize VPA’s effects on the distributed neural circuits underlying reward and energy balance, shifting the setpoint towards increased energy consumption *and* expenditure.

We were surprised to observe that at a dose deemed to be generally “tolerable”, VPA-exposed breeder pairs required at least two weeks longer to produce a litter of pups. Multiple mutually nonexclusive mechanisms relating to male and female factor may explain this subfertility^57^, including endocrine imbalances that impact gonadal health as well as reductions in libido/sexual drive, that may be congruent with deficits in sucrose preference. VPA-exposed pups were generally normal in appearance and by P15 (but not earlier), displayed reduced exploratory behavior on our pup open field assay, without overt evidence of neurological motor impairment. When examined as adults, hourly or total daily distances were unchanged by prenatal VPA exposure. However, VPA-exposed mice displayed more frequent active states that were significantly shorter in duration, illustrating a fundamental derangement in the temporal organization of activity, and phenocopying the effects of social isolation (Fig. 1). In addition to these finer kinematic changes, VPA-exposed mice on average displayed increased sheltering at the cost of feeding, a theme (or “mood”) that we have previously identified in mice modeling Dravet Syndrome and in mice subjected to daily PTZ-induced seizures^32^. Having access to a running wheel was sufficient to overcome their proclivity to shelter, with VPA-exposed mice displaying significantly greater rates of nocturnal wheel-running. In the context of prenatal VPA exposure, we conjecture that increased wheel-running may relate to a tendency to engage in repetitive stereotyped behaviors.

Reductions in social interest, assayed through measures of social play in rats or social exploration in mice, have made the prenatal VPA protocol a widely popular model of autism^25, 26, 74^. The vast majority of these efforts have employed out-of-cage assessments of sociability, such as the three-chamber task^23^. Recently, more continuous home-cage based assessments of social behavior have employed implanted radio-frequency identification (RFID) chips^49, 75^. In our experiments, we applied video-tracking to study dyadic behavior. With continuous, automated and unbiased measures of proximity, we observed no evidence of social withdrawal during their initial interaction. Further, circadian variations in mean proximity were largely similar^49^. During the light spot test^32, 42^, VPA-exposed dyads displayed more prolonged social bouts, revealing if anything, a tendency to *seek* a social partner during an aversive or stressful stimulus. Together, these results call into question the validity of short, subjectively scored assays of social exploration and suggest that broad inferences about “impaired sociability”^76^ may not necessarily translate to truly pervasive changes in social behavior^49, 75^, at least within dyads.

We concede two main limitations to this work. First, by applying VPA dissolved in drinking water, our intention was to avoid the stress of daily (or multiple daily^77^) intraperitoneal injections, which are not without risk when applied to a gravid mouse. Day time reductions in fluid intake^32^ may have resulted in low day time VPA serum levels^39^. VPA’s “carry-over” anticonvulsant effects during the daytime PTZ challenge may be explained by more pervasive increases in brain GABA content, which have been previously shown to more accurately correlate with VPA-induced increases in convulsive threshold^33, 39^. Mini-osmotic pumps (to deliver VPA cerebroventricularly) or subcutaneously absorbed pellets would have been alternative approaches to provide continuous VPA exposure selectively to the female breeder. However, these are invasive options that may themselves impede mating and are not straightforward to terminate (e.g., at parturition). Further, by escaping first pass metabolism, these approaches may result in an altered pattern of metabolites, many of which are known to be biologically active and subserve anticonvulsant roles^33, 39, 45, 57^.

Second, to maximize translational potential, epileptic mice (with spontaneous seizures) would have made for more ideal subjects^78^. These are not a unique entity: many epileptic mouse models are currently available and may each have unique home-cage behavioral signatures. For example, mice with a Dravet syndrome mutation display nocturnal hypoactivity and increased sleep^32^, while nocturnal *hype*ractivity is observed in mice modeling temporal lobe epilepsy^79^. Ideally, to examine how behavioral side effects correlate with anti-seizure efficacy, such studies would be performed with simultaneous electroencephalographic (EEG) monitoring, which may itself impose changes in cage exploration and neurovegetative function. Therefore, as a first pass, we examined VPA’s effects in nonepileptic mice. With improvements in the ergonomics of wireless EEG, we envision a future where preclinical assessments of tolerability will be conducted in an individualized fashion, at doses that are carefully titrated to the cessation of spontaneous seizures.

In conclusion, this work presents a novel paradigm to assess psychotropic and cognitive/behavioral teratogenic side effects of AEDs in mice. Our approach may be extended to any candidate/pipeline neurotherapeutic that would necessitate daily intake and may guide initial safety recommendations regarding ingestion during pregnancy for agents with limited human safety data.

## Acknowledgements

VK receives research support from the NIH (1K08NS110924-01), an American Epilepsy Society Junior Investigator Award (2020), and seed funding from Baylor College of Medicine’s Office of Research.

## Author Contributions

VK conceived and designed the study and drafted the manuscript. JSB and VK designed the figures. JSB, AT, LT, MJJ, PPK and CS contributed to data acquisition and analysis.

## Supplemental Data

**FIGURE S1:**
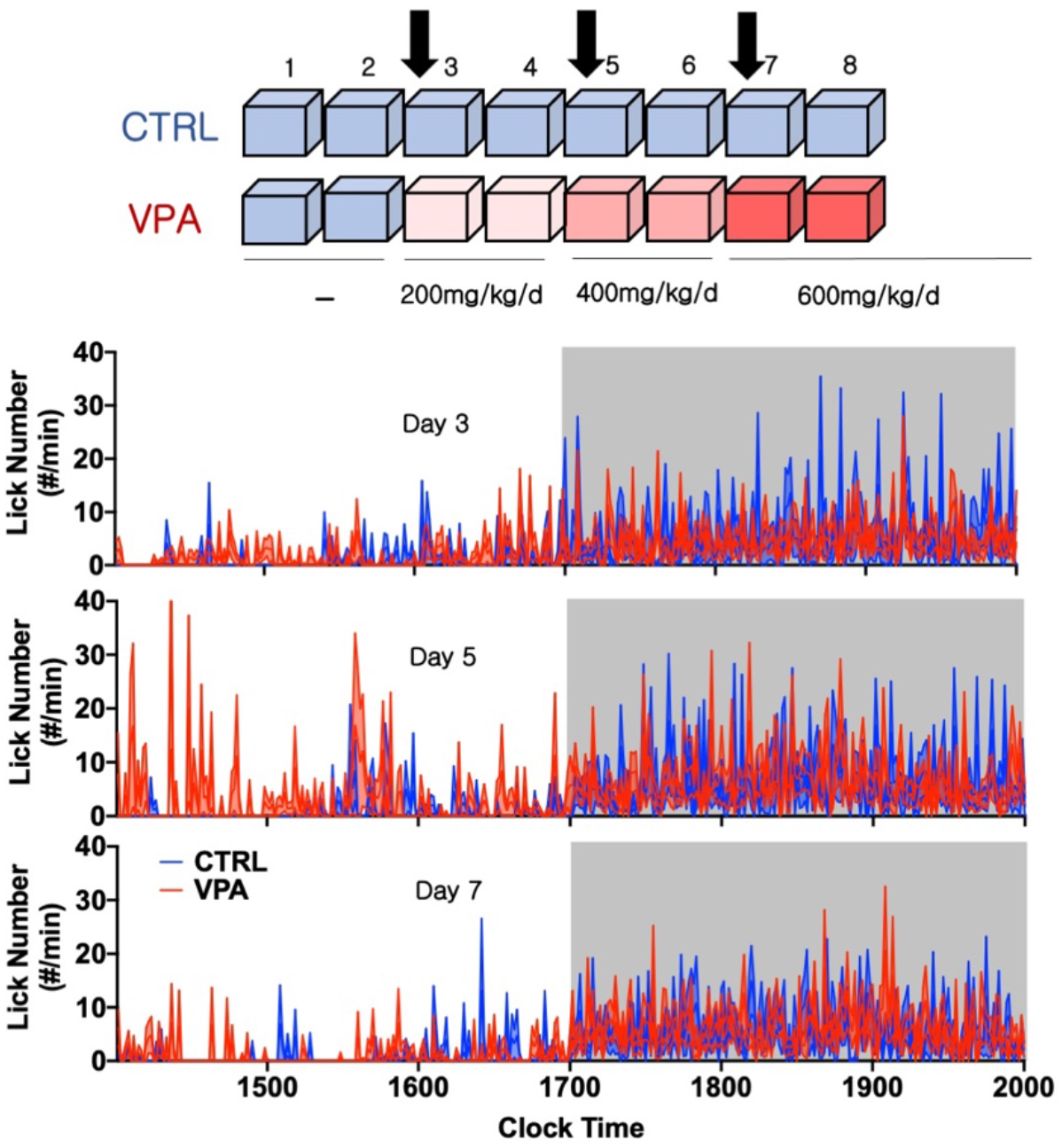
In Fig.1, drinking solutions were replaced for all mice between 1300-1400 on days 3, 5 and 7. CTRL mice received a fresh solution of 0.8% sucrose-drinking water, while VPA-treated mice received fresh solutions with increasing concentrations of VPA. Stepwise increases in VPA concentration did not produce an obvious impact on licking behavior. Mean ± S.E.M shown for all. CTRL: control, VPA: valproic acid.

**FIGURE S2:**
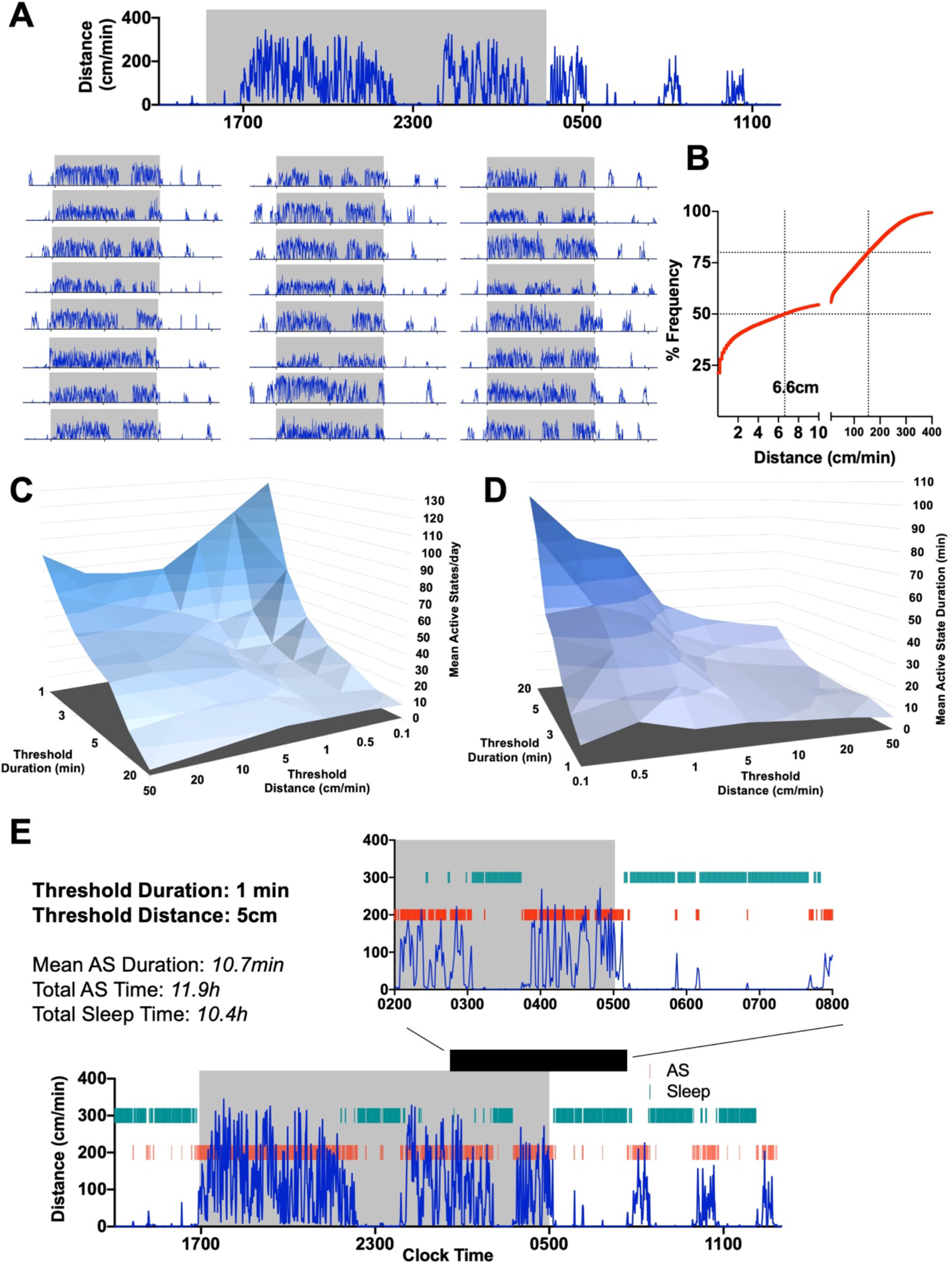
Active state thresholds. (A) Actograms for all C57BL/6J mice [n=25, 12 female] from day 2 (Fig. 1). (B) Cumulative distribution histogram of minutely distance measures, ranging from 0-400cm/min, with 50^th^ percentile corresponding to ~6.6cm. (C) 3-dimensional graph depicting how changes in threshold duration and distance modulate total daily active state numbers, and (D) mean active state duration. Larger duration and distance thresholds effectively ignore short active states. (E) Depiction of active states (AS, red) for a representative mouse at a threshold duration and distance of 1 minute and 5cm/min respectively. Epochs of behaviorally defined “sleep” are shown in green.

**Supplemental Table 1.**

| **Figure** | **Fixed effects (type III)** | **P value** | **P value summary** | **F (DFn, DFd)** |
| --- | --- | --- | --- | --- |
| 1B<br>(Distance) | Day (3-8) | <0.0001 | **** | F (2.462, 56.62) = 10.16 |
|  | Treatment | 0.5684 | ns | F (1, 23) = 0.3349 |
|  | Day x Treatment | 0.9149 | ns | F (5, 115) = 0.2949 |
| 1B<br>(Sleep) | Day (3-8) | <0.0001 | **** | F (2.829, 65.08) = 10.76 |
|  | Treatment | 0.3266 | ns | F (1, 23) = 1.005 |
|  | Day x Treatment | 0.5287 | ns | F (5, 115) = 0.8332 |
| 1B<br>(Sheltering) | Day (3-8) | 0.0004 | *** | F (3.207, 73.76) = 6.615 |
|  | Treatment | 0.9556 | ns | F (1, 23) = 0.003164 |
|  | Day x Treatment | 0.6122 | ns | F (5, 115) = 0.7167 |
| 1B<br>(Feeding Duration) | Day (3-8) | 0.3567 | ns | F (3.437, 80.42) = 1.104 |
|  | Treatment | 0.9418 | ns | F (1, 24) = 0.005446 |
|  | Day x Treatment | 0.071 | ns | F (5, 117) = 2.094 |
| 1B<br>(Lick Duration) | Day (3-8) | 0.116 | ns | F (2.998, 67.16) = 2.044 |
|  | Treatment | 0.2358 | ns | F (1, 23) = 1.482 |
|  | Day x Treatment | 0.1152 | ns | F (5, 112) = 1.817 |
| 1D<br>(Distance) | Time (h) | <0.0001 | **** | F (7.461, 171.6) = 48.01 |
|  | Treatment | 0.5886 | ns | F (1, 23) = 0.3009 |
|  | Time x Treatment | 0.8401 | ns | F (22, 506) = 0.7013 |
| 1D<br>(Sheltering) | Time (h) | <0.0001 | **** | F (7.710, 177.3) = 44.02 |
|  | Treatment | 0.3136 | ns | F (1, 23) = 1.061 |
|  | Time x Treatment | 0.5058 | ns | F (22, 506) = 0.9668 |
| 1E<br>(Active State Number) | Day (3-8) | 0.0007 | *** | F (3.127, 71.91) = 6.238 |
|  | Treatment | 0.1177 | ns | F (1, 23) = 2.641 |
|  | Day x Treatment | 0.5268 | ns | F (5, 115) = 0.8359 |
| 1E<br>(Active State Duration) | Day (3-8) | <0.0001 | **** | F (2.485, 57.15) = 9.916 |
|  | Treatment | 0.8599 | ns | F (1, 23) = 0.03187 |
|  | Day x Treatment | 0.4176 | ns | F (5, 115) = 1.006 |
| 1E<br>(Total AS Time) | Day (3-8) | <0.0001 | **** | F (3.682, 84.68) = 12.62 |
|  | Treatment | 0.1531 | ns | F (1, 23) = 2.183 |
|  | Day x Treatment | 0.1009 | ns | F (5, 115) = 1.893 |
| 1G<br>(Wheel Rotations) | Time (h) | <0.0001 | **** | F (4.657, 102.5) = 20.23 |
|  | Treatment | 0.5632 | ns | F (1, 22) = 0.3445 |
|  | Time x Treatment | 0.6411 | ns | F (21, 462) = 0.8622 |
| 2C<br>(Body Weight) | Week | <0.0001 | **** | F (1, 39) = 80.91 |
|  | Treatment | 0.1631 | ns | F (1, 39) = 2.021 |
|  | Week x Treatment | 0.5696 | ns | F (1, 39) = 0.3289 |
| 2D<br>(Distance) | Time (h) | <0.0001 | **** | F (8.689, 338.9) = 70.11 |
|  | Treatment | 0.9807 | ns | F (1, 39) = 0.0005927 |
|  | Time x Treatment | 0.0068 | ** | F (22, 858) = 1.920 |
| 2D<br>(Sheltering) | Time (h) | <0.0001 | **** | F (8.655, 337.5) = 81.12 |
|  | Treatment | 0.1107 | ns | F (1, 39) = 2.664 |
|  | Time x Treatment | 0.3069 | ns | F (22, 858) = 1.130 |
| 2D<br>(Feeding Entries) | Time (h) | <0.0001 | **** | F (6.161, 240.3) = 41.46 |
|  | Treatment | 0.172 | ns | F (1, 39) = 1.936 |
|  | Time x Treatment | 0.0309 | * | F (22, 858) = 1.648 |
| 2J<br>(Wheel Rotations) | Time (h) | <0.0001 | **** | F (3.648, 138.6) = 16.60 |
|  | Treatment | 0.0027 | ** | F (1, 38) = 10.34 |
|  | Time x Treatment | <0.0001 | **** | F (20, 760) = 3.569 |
| 4C<br>(Distance) | Time (h) | <0.0001 | **** | F (9.459, 539.2) = 74.80 |
|  | Treatment | 0.7746 | ns | F (1, 57) = 0.08279 |
|  | Time x Treatment | 0.3861 | ns | F (22, 1254) = 1.060 |
| 4C<br>(Sheltering) | Time (h) | <0.0001 | **** | F (11.17, 636.4) = 83.41 |
|  | Treatment | 0.0141 | * | F (1, 57) = 6.417 |
|  | Time x Treatment | 0.2649 | ns | F (22, 1254) = 1.171 |
| 4C<br>(Feeding Entries) | Time (h) | <0.0001 | **** | F (8.606, 481.9) = 59.05 |
|  | Treatment | 0.1576 | ns | F (1, 56) = 2.052 |
|  | Time x Treatment | 0.0386 | * | F (22, 1232) = 1.601 |
| 4G<br>(Wheel Rotations) | Time (h) | <0.0001 | **** | F (5.097, 262.8) = 30.89 |
|  | Treatment | 0.0476 | * | F (1, 53) = 4.113 |
|  | Time x Treatment | 0.0022 | ** | F (20, 1031) = 2.164 |
| 5C<br>(Sum Distance) | Time (h) | <0.0001 | **** | F (7.490, 172.3) = 46.30 |
|  | Treatment | 0.2372 | ns | F (1, 23) = 1.473 |
|  | Time x Treatment | 0.0063 | ** | F (22, 506) = 1.948 |
| 5C<br>(Distance Between Subjects) | Time (h) | <0.0001 | **** | F (9.766, 222.8) = 39.40 |
|  | Treatment | 0.2485 | ns | F (1, 23) = 1.402 |
|  | Time x Treatment | 0.0072 | ** | F (22, 502) = 1.926 |

